# Doxycycline Significantly Enhances Induction of iPSCs to Endoderm by Enhancing survival via AKT Phosphorylation

**DOI:** 10.1101/2020.04.13.034595

**Authors:** Cristina Esteva-Font, Tao Su, Caitlin Peaslee, Caroline Duwaerts, Ke Liu, Marisa Medina, Jacquelyn J. Maher, Aras N. Mattis

## Abstract

Induced pluripotent stem cells (iPSCs) provide an important tool for the generation patient-derived cells including hepatocyte-like cells via developmental cues through an endoderm intermediate. However, most iPSCs fail to differentiate into endoderm, with induction resulting in apoptosis. To address this issue, we built upon published methods to develop an improved protocol with our discovery that doxycycline dramatically enhances the iPSC to endoderm differentiation efficiency by inhibiting apoptosis and promoting proliferation via the AKT pathway. We tested this new protocol in more than 70 iPSC lines with consistent formation of complete sheets of endoderm in 90%. Endoderm generated by our method achieves similar transcriptomic profiles, including FOXA2, HNF1β, CXCR4, and SOX17 positive cells, and the ability to be further differentiated. Furthermore this method achieves a four-fold increase in endoderm cell number and will accelerate studies of human diseases *in vitro* and facilitate the expansion of iPSC-derived cells for transplantation studies.

## Introduction

IPSC-derived hepatocytes (iPSC-Heps) effectively model single nucleotide polymorphism (SNP) based inter-human metabolic differences including cytochrome P450 CYP2D6 type I and type II metabolic differences (Takayama et al., 2014). More importantly, unlike hepatoma tumor lines, iPSC-Heps function as post-mitotically differentiated cells without significant proliferation, but with the added benefit of true high-level hepatic functioning. Unlike primary human hepatocytes, we can readily expand iPSCs to differentiate into any number of hepatocytes and can genetically modify them in forward genetic screening. Thus iPSC-Heps have clear potential to be the most useful individual human-specific cell line to recapitulate not only our variable differences in SNP gene expression but also as a potential way to model our individual increased or decreased susceptibility to disease. Given the promise of iPSC-Heps, their Achilles heel has been the tremendous variability encountered during efforts to differentiate them into hepatocytes. While embryonic stem cell lines show greater reliability in differentiation, different iPSC lines, even if derived from a single patient, often succumb to apoptosis before reaching the endoderm stage. IPSC lines that survive to endoderm typically go on to successfully produce hepatocytes.

In the published literature and in practice, research groups often employ only several highly-selected subclones of pluripotent cells for all of their studies, because they lack efficient induction of endoderm. In order to usefully characterize patient-derived iPSC lines from multiple patients using high content screens, it is critical to consistently make true endoderm in complete monolayer sheets. We therefore explored the ability to achieve more efficient induction of endoderm.

Differentiation of stem cells into mature cell lines often progresses through a step-wise natural progression similar to embryogenesis. This differentiation progresses through the three germ layers, ectoderm, endoderm and mesoderm. Of these, for unknown reasons, production of endoderm appears to be particularly challenging in vitro and perhaps this is why spontaneous mature teratomas in vivo often contain more ectodermal and mesendodermal tissues than endoderm (Sahin et al., 2017). Yet if the promise of iPSCs is to produce mature cell lines from any patient or healthy individual, this issue must be overcome. Induction of human iPSCs to endoderm via Activin A often produces significant apoptotic cell death. High dose Activin A induces the NOGGIN pathway driving multiple pathways important in endoderm induction including SMAD 2/3/4 activation. Unfortunately, there are also multiple downstream pathways activated by Activin A driving cellular apoptosis including SHIP-1, DAP-Kinase, and TIEG (downstream of SMAD signaling), and inhibition of AKT pathways (via inhibition of PI3KINASE) (Charlier et al., 2010; Wang et al., 2002; Zhao et al., 2018a). Because these apoptotic pathways provoke significant loss of cells, successful induction is dependent on health of the cell lines, plating density, media balance, and surface environment-making differentiation to endoderm extremely challenging. Even under optimal conditions, most iPSC lines undergo total apoptosis at the endoderm induction stage within the first two to three days. In fact, multiple investigators have categorized cell lines as either good or incompetent to form endoderm. Our hypothesis was that the problem was a technical hurdle because of the inappropriate Activin A activation of apoptotic pathways via SMAD signaling, and that there must be an efficient way to circumvent apoptosis and induce any cell line to efficiently form endoderm.

We therefore set out to test endoderm induction via genetic methods. In the process we discovered that the use of doxycycline as a medium additive significantly enhanced endoderm formation and subsequent differentiation. We therefore fully explored the use of doxycycline to induce endoderm and finally differentiate our cells to iPSC-Heps. Our data show that doxycycline significantly improves survival and expansion at the endoderm stage, and thus improves differentiation of iPSCs from multiple donors and different reprograming conditions into sheets of endoderm cells with high efficiency. Our results also illustrate that there are no significant differences in gene expression between iPSC-Heps generated with or without doxycycline. One benefit of doxycycline is its ability to stimulate phosphorylation of AKT, supporting survival. Our protocol induces a strong proliferative response at the endoderm stage.

## Results

### Induction of multiple iPSC lines from different patients using defined media

With our interest in characterizing multiple different iPSC-Heps, we tested and found significant differentiation variability and cell death during endoderm induction using popular published protocols (Ang et al., 2018; Ma et al., 2013; Si-Tayeb et al., 2010). In order to better optimize our system, we selected one Matrigel lot for all experiments based on its ability to somewhat support our induction of our weakest iPSC lines; however, even with this Matrigel lot, we did not see efficient differentiation in such lines (**Figure 1C**).

**Figure 1.**
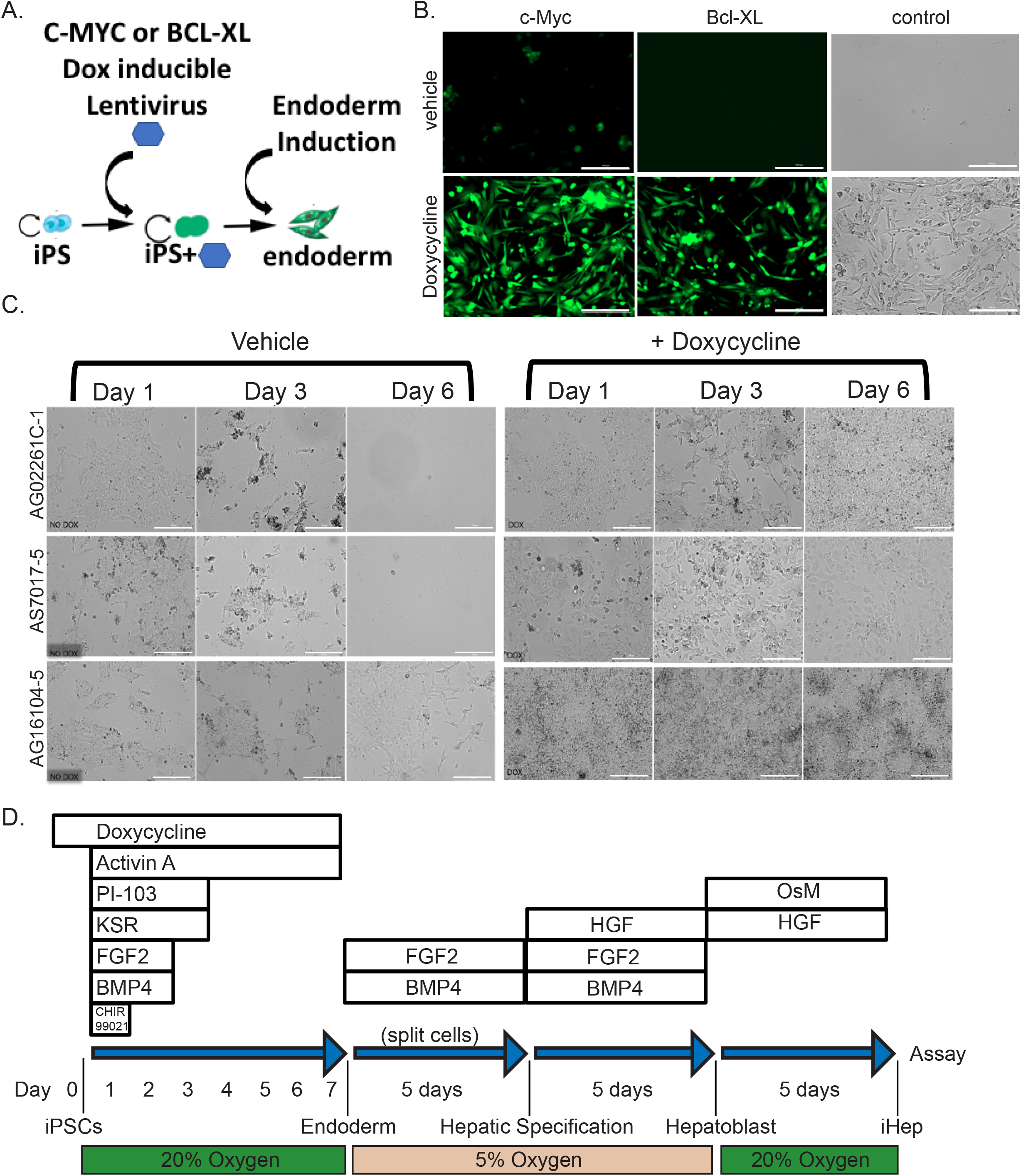
Doxycycline Rescues Induction to Endoderm. (A) Schematic of C-MYC or BCL-XL doxycycline inducible transduction and establishment of iPSC lines that were then able to efficiently form definitive endoderm. (B) Immunofluorescence and bright-field images showing induction of definitive endoderm from iPSCs after 6 days using protocol shown in S. Figure 1A, with and without doxycycline in iPSCs with established doxycycline-inducible C-MYC, BCL-XL or control (no viral transduction) in cell line AS7017-5 (scale bar = 100μM). Control experiments without virus rescued induction of cell line AS7017-5 (n=2). (C) Bright-field images from three inefficient iPSC lines induced with or without the addition of doxycycline (no additional transgenes), scale bar = 100μM, (n=3). (D) Schematic for optimal induction of iPSCs into definitive endoderm (days 0-7) and then to iPSC-Heps. Protocol also referred to as ML1.

We next tested combining different protocols to see if we could achieve better differentiation across multiple independent cell lines. Addition of CHIR99021 in the first 24 hours improved the number of cells in multiple different cell lines (Rashid et al., 2010). Addition of FGF2 and BMP4 for the first 48 hours sometimes, but not always, helped endoderm induction (Rashid et al., 2010). We chose to keep these factors in because there was improvement in some lines (Xu et al., 2011). Based on our prior experience with induction in serum free media (Ma et al., 2013; McLean et al., 2007), we kept serum out of many early experiments. However, in frustration with very fragile lines, we finally added knockout serum replacement (KSR) and found that addition of small amounts of KSR in the first 72 hours improved survival. In order to counteract the insulin present in KSR, we added the small molecule PI-103 (PI3 kinase inhibitor) (Loh et al., 2014; McLean et al., 2007). This combined protocol (**Supplementary Figure 1A**), worked for about one fourth of our cell lines but still showed stochastic cell death, inconsistency, and still did not allow us to differentiate numerous important patient-derived iPSCs.

### Doxycycline rescues endoderm induction independent of transgenes

We explored the possibility to artificially turn on proliferation while inhibiting apoptosis, to short-circuit cell death and force endoderm induction in our cell lines. We were inspired by Hirose et al. that were able to improve erythroblast differentiation via induction of BCL-XL and cMYC (to inhibit apoptosis and activate cellular proliferation respectively) for the improved production of erythrocytes from iPSCs (Hirose et al., 2013). We wanted to use a similar strategy for our induction of endoderm. We chose poorly differentiating iPSC lines (AG02261C-1, AS7017-5, AG16104-5; **Figure 1**), and infected them with green fluorescent protein (GFP)-tagged lentivirus expressing Dox-On BCL-XL or Dox-On cMYC (**Figures 1A and 1B**). We isolated clonal GFP-positive iPSC subclones by fluorescence-activated cell sorting (FACS) and expanded pure populations of these iPSCs containing integrated BCL-XL or cMYC. Next we tested endoderm induction, by providing doxycycline inducing BCL-XL or cMYC (**Figure 1B**). Turning on BCL-XL or cMYC immediately rescued these inductions from cellular death. Using this system, we were able to differentiate three poorly differentiating cell lines when either cMYC or BCL-XL was induced by doxycycline (**Figure 1B**). BCL-XL and cMYC inductions were confirmed by quantitative PCR and endoderm was confirmed by immunohistochemical staining (**S.Figures 1B and 1D**).

Surprisingly, we noted that control iPSC lines not transduced with lentivirus but treated with doxycycline also survived induction to iPSC-derived endoderm (**Figure 1B**). We confirmed this result using multiple independent poorly-differentiating cell lines (**Figure 1C**). With this discovery, we abandoned the use of lentivirally transduced iPSCs and focused on characterizing the effects of doxycycline on iPSC-derived endoderm. To further analyze the ability of doxycycline to induce endoderm, we titrated iPSC seeding density versus doxycycline doses on five different lines with variable inducibility from completely uninducible to inducible without doxycycline (**S. Figure 1C**). At even the lowest doses of doxycycline, there was an immediate improvement in most lines. Most were able to induce to confluency and avoided cell death. The data show that addition of even small concentrations of doxycycline across a variety of plated cell densities was able to improve nearly all of the iPSC lines tested for endoderm induction and even allowed us to form complete monolayers in many lines in 96-well format. Because doxycycline is an antibiotic and because it is known to inhibit mitochondrial translation, we tested other antibiotic classes including ampicillin, ciprofloxacin, gentamycin, and puromycin. Compared to doxycycline, only gentamicin showed a similar statistically significant ability to enhance survival of cells during induction to endoderm but with altered kinetics at equivalent doses (**Figure 2B**). Because there were reports of both doxycycline and gentamycin affecting mitochondrial calcium channels (Sastrasinh et al., 1982; Schwartz et al., 2013), we tested inhibitors CGP-37157, KB-R7943 and ruthenium red that could inhibit mitochondrial sodium and/or calcium exchanger as well as uptake and release from mitochondria. These compounds were not nearly as effective as doxycycline in two separate lines tested (**Figure 2B** and **S. Figure 4**).

**Figure 2.**
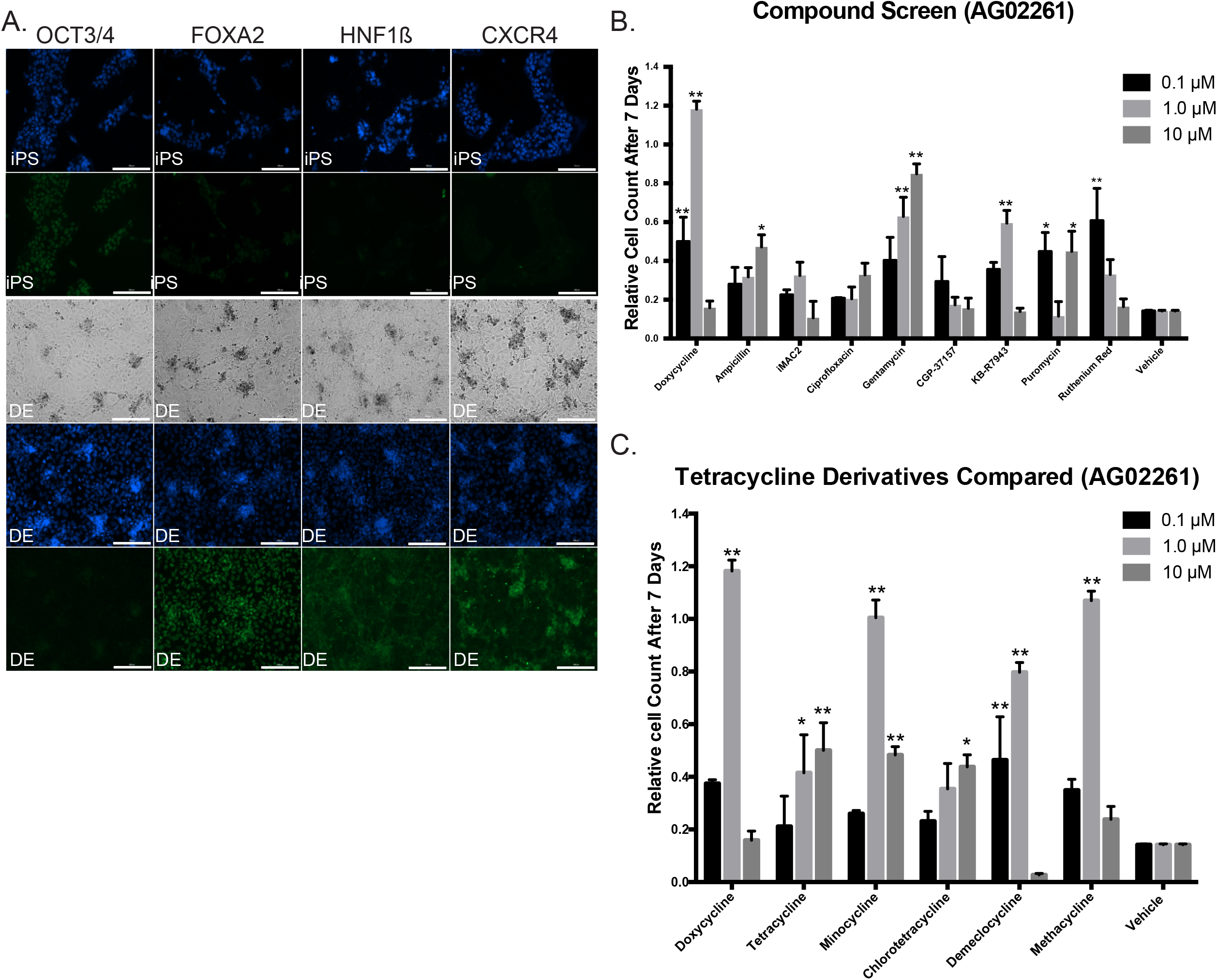
Doxycycline Enhanced Endoderm and Testing of Additional Compounds. (A) Immunofluorescence and bright-field images showing induction of iPSC line 1023-5 to definitive endoderm (DE) with Hoechst 33342 nuclear staining in blue, and antibody staining for OCT3/4, FOXA2, HNF1β and CXCR4 with AlexaFluor 488 secondary (green) antibody. Bright-field microscopy shows complete monolayer. (scale bar = 200μM). (B) Compound Screen for iPSC induction to endoderm using other antibiotics and other compounds affecting the mitochondria compared to doxycycline as additives to the protocol. Compounds were added instead of doxycycline to protocol ML1. While most antibiotics failed to have the same effect, surprisingly Gentamycin was shown to also have a strong increasing dose response. KB-R7943 and Ruthenium Red also showed responses at specific doses in one cell line shown here (AG02261) but not another cell line (AS7192) in S. Figure 4B. The number of cells is shown measured seven days after induction compared to the plating density at day 0 (n=3 technical replicates for each independent cell line). (C) Comparison of Tetracycline Derivatives in iPSC lines AG02261. Doxycycline was compared to tetracycline derivatives in its ability to rescue endoderm induction. Minocycline, and Methacycline showed similar ability to rescue endoderm at the 1 μM concentration with toxicity at the 10μM concentration. Demeclocycline showed a similar trend. The number of cells is shown measured seven days after induction compared to the plating density at day 0 (n=3 technical replicates for each independent cell line). Data analyzed by Two way ANOVA and presented as mean ± SEM. * p < 0.05; ** p < 0.01.

We also tested tetracycline derivates for their ability to rescue induction to endoderm, and compared to doxycycline, only minocycline, demeclocycline and methacycline showed similar trends in two different cell lines. The parent compound tetracycline and derivative chlorotetracycline did not appear to work as effectively (**Figure 2C** and **S. Figure 4A**).

We next tested the effects of doxycycline on iPSC growth and found that it was able to improve daily growth kinetics in nearly all lines tested (**S. Figure 2A**). A dose-response study demonstrated a beneficial effect of doxycycline up to 3μg/mL but eventually toxicity at higher levels (**S. Figure 2B**). We validated induction of iPSCs to endoderm in our protocol by immunohistochemical staining for markers of endoderm including FOXA2, HNF1β, and CXCR4 (**Figure 2A**). Our protocol, for convenience designated ML1 (**Figure 1D**), showed efficient induction of endoderm. More importantly the ML1 protocol allowed induction of complete monolayers of endoderm from nearly all iPSC lines tested, even those that failed other protocols recommended for poorly differentiating iPSC (Ang et al., 2018; Si-Tayeb et al., 2010).

### Early reduction of cleaved-caspase 3 cleavage

As a first step to understand the mechanism by which doxycycline was affecting induction of iPSCs to endoderm, we explored whether doxycycline was affecting cell proliferation, cell death, or both. For this, we performed an induction for endoderm using protocol ML1 for 24 hours and then fixed and stained the cells for cleaved caspase 3 (CC3) and the proliferation marker Ki-67. After quantification, we noted both significant Ki-67 staining and cleaved caspase 3 (CC3) staining as expected (**Figures 3A and S. Figure 3A**). In the presence of doxycycline, during this early induction timepoint, we noted a greater than 50% decrease in CC3 staining with only a very slight decrease in overall Ki-67 levels over 4 different cell-line experiments. We concluded that this early reduction in apoptosis leads to sustained proliferation leading to successful induction.

**Figure 3.**
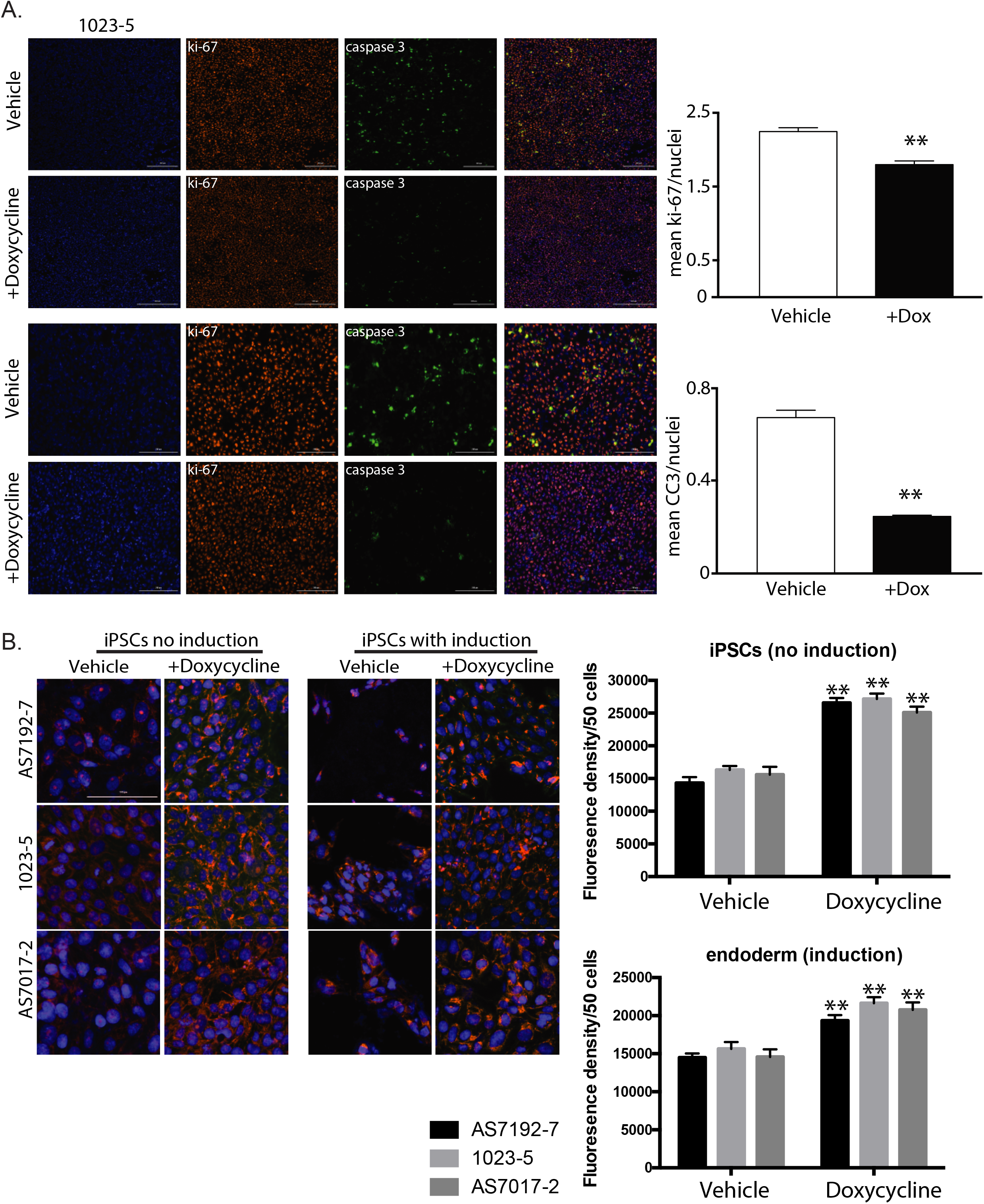
Doxycycline Reduces Apoptosis and Increases Mitochondrial Mass. (A) Immunofluorescence of iPSC line 1023-5 on day 2 of induction for Hoechst 33342, Ki-67, Cleaved Caspase 3 (CC3), and combined shows greatly decreased CC3 staining with doxycycline treatment. At this early time point, there is a small but significant decrease in Ki-67. First two rows are 10x with scale bar at 400μm and bottom two rows 20x with 200μm scale bar. (B) Mitotracker Orange staining of iPSCs with no induction and with induction shows that addition of Doxycycline increases the quantity of polarized mitochondria compared to vehicle alone in three separate iPSC lines (AS7192-7, 1023-5, and AS7017-2) (n=3). Scale bar represents 100μm. Data were analyzed by Student’s t test and presented as mean ± SEM. * p < 0.05; ** p < 0.01.

### Interrogation of mechanism reveals multiple pathway interactions

Given its function as an antibiotic, early studies of doxycycline identified it as a mitochondrial translational inhibitor (Chatzispyrou et al., 2015). However, newer studies have shown additional functions. For example, in some tumor lines doxycycline can alter numerous metabolic genes (Ahler et al., 2013; Lamb et al., 2015), with even long-term epigenetic modifications noted in T-cells after treatment (Becker et al., 2016).

We hypothesized that TGF-β-activated differentiation protocols via Activin A under these culture conditions limit cell division, because under certain cellular scenarios activates the apoptotic machinery. We speculated that perhaps doxycycline by inhibiting mitochondrial function, was interfering with the ability of the mitochondrial apoptosis machinery to fully engage. Therefore, to characterize mitochondria, we performed Mitotracker staining with and without doxycycline on iPSCs with and without induction to endoderm (**Figure 3B**). Treatment of both iPSCs and induced cells with doxycycline showed increased MitoTracker Orange CMTMRos staining, consistent with an increase of functional mitochondrial mass. Doxycycline therefore does not affect mitochondria in iPSCs and induced iPSCs in the same way that it does in cancer stem cells (Lamb et al., 2015; Yang et al., 2015; Zhong et al., 2017).

We further sought to delineate the pathways that were leading to increased apoptosis during the endoderm inductive phase. Induction of endoderm by high-dose activin A is thought to drive the nodal pathways by binding Activin A receptors 1 and 2, thus driving the phosphorylation of SMAD2 and 3 (Yu et al., 2015). The Phospho-SMAD2/3 complex recruits SMAD4 and enters the nucleus to drive transcription of multiple downstream targets (Yu et al., 2015). There are several published mechanisms for activated SMAD2/3/4 connection to the apoptotic pathway. First, TGFβ is known to induce apoptosis through SMAD-mediated DAP-kinase expression (Jang et al., 2002). Secondly, TGFβ receptor (Activin A activated) is also directly connected to a SMAD4 mitochondrial translocation and cyctochrome c oxidase subunit II interaction (Pang et al., 2011). These findings might explain the apoptotic propensity of iPSCs from certain cell lines during endoderm differentiation.

In order to more definitely identify mechanisms involved, we attempted to induce and differentiate eight iPSCs with and without doxycycline in pairs. Of the eight iPSCs without doxycycline, we were able to generate sufficient endoderm for five out of the eight, and sufficient iPSC-Heps in four of eight samples to extract quality RNA for RNA Sequencing analysis (**S. Figure 5B**). In contrast, using doxycycline, we were able to easily generate enough endoderm and iPSC-Heps for RNA Sequencing from all eight of the samples. We performed RNA Sequencing with the samples that passed quality control tests. After blinded clustering analysis, we noted that all endoderm, iPSC-Heps, and IPSCs clustered appropriately in our heatmap without any bias for treatment with or without doxycycline (**Figure 4A**).

**Figure 4.**
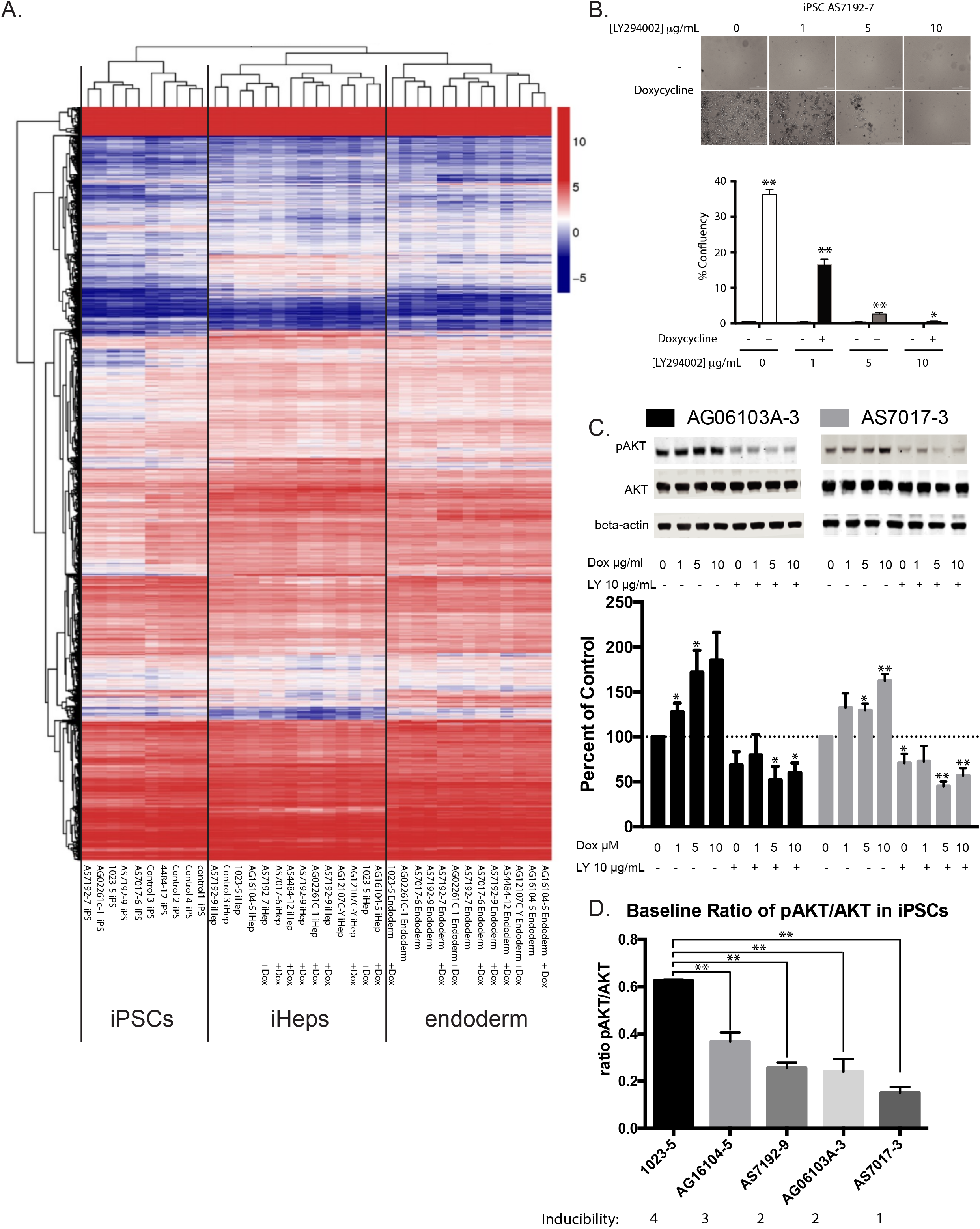
Mechanism of Doxycycline Effects on Definitive Endoderm Induction. (A) RNA Sequencing of iPSC, definitive endoderm, and final iPSC-Heps cluster appropriately after unbiased hierarchical clustering analysis with and without doxycycline enhanced differentiation during the endoderm phase. (B) Doxycycline enhanced definitive endoderm induction can be inhibited by LY294002 in a dose-dependent way (iPSC line AS7192-7 shown) (n=3 technical replicates in 3 different cell lines). (C) Escalating doses of doxycycline cause increased P-AKT/AKT ratio. LY294002 inhibited P-AKT levels are not reversible by increasing doses of doxycycline (iPSC lines AG06103A-3 and AS7017-3 shown) (n=3 technical replicates in 3 different cell lines). Western blots quantified with Odyssey LiCor imaging system. (D) Pre-existing baseline ratio levels of pAKT/AKT in individual iPSC lines correlate with the ability to induce these lines to definitive endoderm (n=3 technical replicates for each cell line). Data were analyzed by Student’s t test versus control and presented as mean ± SEM. * p < 0.05; ** p < 0.01.

Upregulated and downregulated RNA pathways are shown in **Sup. Table 1**. We found 205 genes were significantly downregulated and 47 upregulated (adjusted p value of 0.05). Ingenuity Pathway Analysis (IPA) was performed which revealed Cellular Growth and Proliferation as well as Cell Death and Survival as within top Molecular and Cellular Functions during induction to endoderm. However, these pathways appeared largely balanced without clear targets that were overwhelmingly affected. Notably levels of Caspase 3 were minimally but significantly reduced. In addition, Inhibition of matrix metalloproteases (MMPs), a known target of doxycycline was a top canonical pathway. The connection between MMPs and apoptosis has been explored (Mannello et al., 2005), however, only TIMP4 inhibition would be consistent with decreased apoptosis in this data set.

To further explore affected protein pathways including regulatory protein modifications such as protein phosphorylation that would not be detected by RNA analysis, we performed a quantitative antibody array (Moon Bio, Sunnyvale, CA) on protein extracts from an induced iPSC-endoderm line with and without doxycycline (**Sup. Table 2**). Increased phosphorylation was noted following doxycycline in protein pathways driving proliferation and cell survival including Elk-1 Ser383, ACK1 Tyr284, Dok-2 Tyr299, MEF2C Ser396, and HER2 Tyr1221/Tyr1222 (Frogne et al., 2009; Lavaur et al., 2007; Pon and Marra, 2016; Shinohara et al., 2005; Zhao et al., 2018b). Phosphorylation of ACK1 Tyr284 has been shown to lead to increased AKT phosphorylation (Zhao et al., 2018b).

As a next step, and to confirm that AKT was indeed hyperphosphorylated in the presence of doxycycline, we used four iPSCs and performed a doxycycline dose response experiment. Treatment of iPSCs during induction of endoderm showed protection from cell death as we previously showed which was entirely reversible with increasing doses of LY294002 (**Figure 4 B**). Furthermore, induction at 1, 5, and 10μM doxycycline showed dose-dependent increases in phosphorylated AKT levels which was entirely reversible with LY294002 (**Figure 4C**).

Why different iPSCs show differential sensitivity to Activin A-induced cell death is puzzling. AKT directly interacts with unphosphorylated SMAD3, through a mechanism independent of AKT kinase activity. By sequestering unphosphorylated AKT, the complex can default into apoptosis (Conery et al., 2004). Therefore, pre-existing levels of phosphorylated AKT could determine the efficiency of induction and this may hold the key balance. In order to characterize if such levels could be leading to the noted differences between cell lines, we characterized the inducibility of cell lines as excellent (four stars), average (three stars), and poorly inducible cell lines (2 and 1 stars). We then compared preexisting levels of Phospho-AKT/AKT levels in these iPSCs (**Figure 4D**). Excellent inducers showed baseline P-AKT/AKT ratios of greater than 0.3 versus poor inducers.

### Enhanced endoderm induction does not alter the final hepatocyte maturity

We wanted to confirm that use of doxycycline did not affect the final iPSC-derived hepatocytes. To characterize this, using our RNA Sequencing data on iPSC-Heps (**Figure 4A**), we did not see any statistically significant findings in iPSC-Heps. Doxycycline did not alter the iPSC-Heps (**Figure 4A**).

### Addition of Y-27632 plus doxycycline allowed induction of severely induction resistant cell lines

To determine the value of other medium additives in our differentiation protocol relative to doxycycline, we performed a systematic elimination experiment but in order to achieve efficient differentiation in most of our lines, all of our additives were required (**Figure 5A**). In our extensive attempt to differentiate numerous iPSCs, we did note some rare cell lines that continued to undergo significant apoptosis. Realizing that apoptosis can be driven by additional pathways, such as the Rho kinase pathway (Shi and Wei, 2007), we tested the addition of Y-27632. By itself Y-27632 was not able to rescue resistant lines during induction to iPSC-derived endoderm. Because both Y-27632 and doxycycline are considered to work on different pathways, we tried adding doxycycline and Y-27632 together to poorly differentiating cell lines (**Figure 5B**). Using this modified protocol, we were able to now differentiate 42 iPSCs from **Supplementary Table 3** and an additional 28 iPSC lines from the Maher lab to fully confluent wells for downstream analysis. In addition, given doxycycline’s prior report to help iPSC cell growth (Chang et al., 2014), we also found that addition of doxycycline with Y-27632 helped improve iPSCs survival during single cell FACS sorting (**S. Figure 5B**).

**Figure 5.**
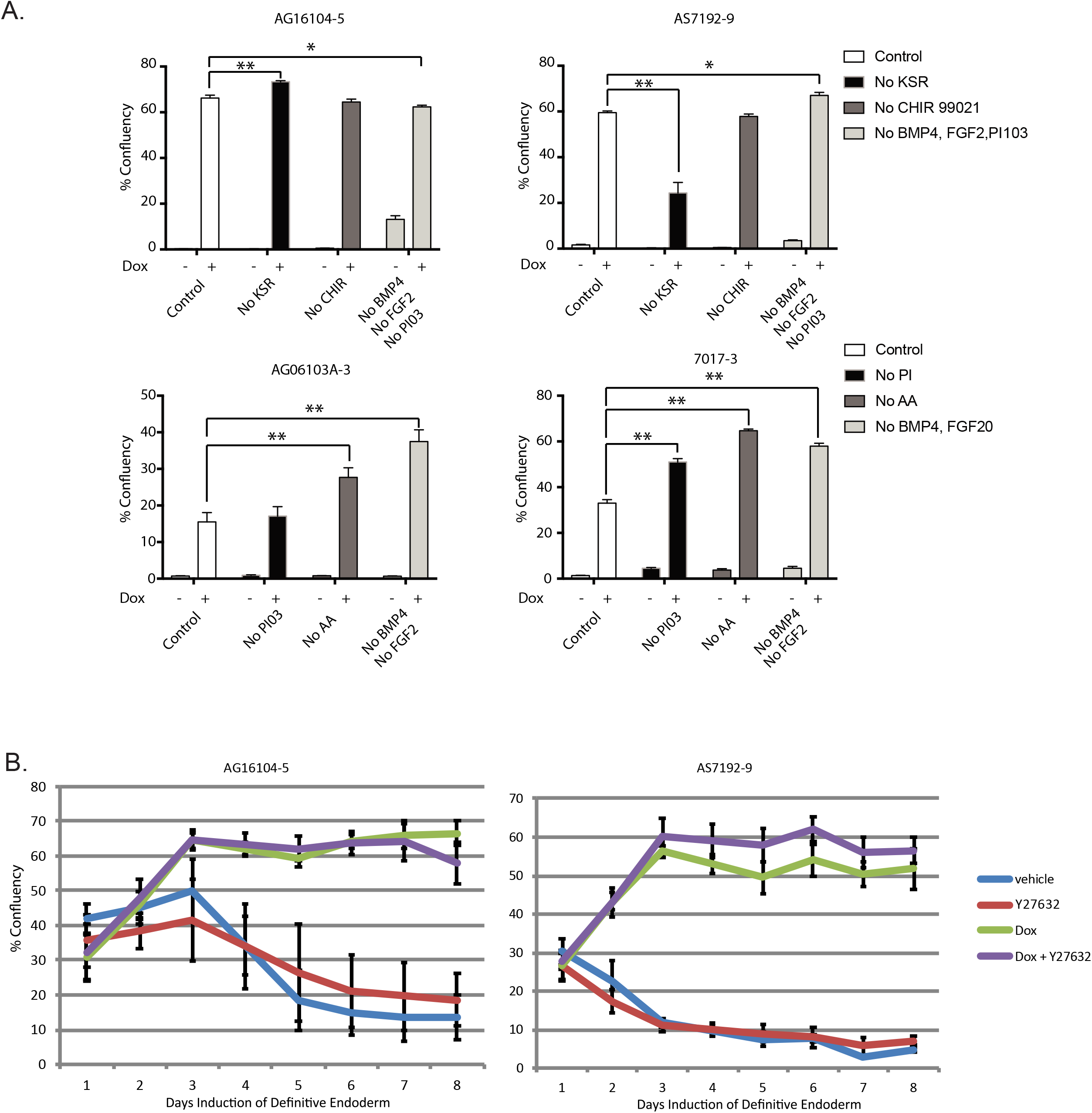
Optimization of ML1 IPSC to endoderm protocol and Difficult cases of Induction. (A) Experiment with regents excluded in an attempt to minimize components within induction media. Our conclusion is that all reagents are required (n=3 technical replicates for each of the 4 cell lines). (B) Percentage well saturation during induction of iPSCs with vehicle (blue line), Y-27632 (red line), Doxycycline 1 μg/mL (green line), or Doxycycline plus Y-27632 (n=3 technical replicates for each cell line). Data analyzed by Two way ANOVA and presented as mean ± SEM. * p < 0.05; ** p < 0.01. Error bars in B represent 95% confidence interval.

## Discussion

Differentiation of iPSCs to hepatocytes has long been a challenging process. Due to Activin A-induced apoptosis, iPSC induction to endoderm is often unreliable; this causes frustrating expenditures of time and resources, and in many cases prevents the completion of critical experiments with patient-derived cell lines. Doxycycline seems to play a dual role in the case of helping endoderm induction. During the early phase, it appears to inhibit apoptosis, while during the late phase it seems to promote proliferation via AKT phosphorylation. Doxycycline was developed from Terramycin, a naturally occurring antibiotic isolated from the actinomycete, *Streptomyces rimosus* in 1950 (Finlay et al., 1950). Much like most natural antibiotics, Terramycin was able to inhibit gram-positive and gram negative cocci. Interestingly the derivative antibiotics in our case including doxycycline, minocycline, and methacycline were able to effectively rescue induction but not the parent compound tetracycline. We also note that some published protocols routinely use antibiotics in differentiation protocols likely to prevent infection but can obviously affect differentiation including survival or death of cells.

Maintenance of pluripotency in iPSCs is generally a metabolic process, and studies have tried to show that during successful endoderm induction, there is a switch from glycolysis to oxidative phosphorylation (Gu et al., 2016; Moussaieff et al., 2015). However, doxycycline is known to inhibit mitochondrial function and therefore reduce oxidative phosphorylation. Inhibition of mitochondrial metabolism and successful induction of endoderm are contrary to the notion that endoderm induction requires an oxidative metabolic switch. On this same idea, it was previously reported that deletion of cMYC leads to spontaneous primitive endoderm differentiation from iPSCs (Smith et al., 2010) cMYC is thought to regulate some of the metabolic switching between glycolysis and oxidative metabolism, however, newer work confirms that differentiation to endoderm does not appear to be affected by MYC re-expression or change of the metabolic switch (Cliff et al., 2017). Our results show that induced cMYC expression is entirely compatible with endoderm induction, without negative effects on net endoderm or final iPSC-Heps differentiation. Clearly there is still much to learn about induction of endoderm.

The mechanism of doxycycline enhancing endoderm induction via AKT phosphorylation is also supported by pathways highlighted by our RNA expression data. For example, we noted that the ERK pathway that induces apoptosis was also affected via downregulation of CAV1 (Liu et al., 2001) and that furthermore CAV1 can also interact with TGF-β receptor influencing SMAD2 phosphorylation (Razani et al., 2001). It is likely that additional pathways are further affected requiring additional study. We also noted that gentamicin rescued endoderm induction in our limited screens and while we tried to find a common mechanism with doxycycline, possibly related to mitochondrial calcium channels, we were not successful and would like to follow up on this result.

The use of this new protocol has revitalized our entire differentiation pipeline and allowed our lab to perform the experiments that were previously out of reach because of the inefficiency of endoderm induction and subsequent hepatocyte differentiation. With the addition of doxycycline, in vitro differentiation of iPSC to hepatocytes is now possible from any patient and almost any line, making phenotype analysis possible and comparison of patients in parallel experiments. We look forward to additional modifications to the protocol enabling complete maturation of iPSC hepatocytes to fully mature hepatocytes from each of the zonal areas of the liver lobe.

Here we show that doxycycline is an effective and compatible compound, for the induction of iPSCs to endoderm that significantly enhances survival and increases the number of endoderm cells by at least four fold. Given this increased proliferation of cells at the endoderm stage, this method produces cells with nearly equivalent hepatocyte-like gene expression with more consistency after final differentiation. This has significant implications for cellular therapy but also highlights the importance to understand doxycycline effects on cellular metabolism and alternative pathways when used in models of gene regulation.

## Experimental Procedures

### Cell Culture of iPSCs

The media conditions to maintain human iPSCs have all been previously described (Ludwig et al., 2006). Cells were grown on growth factor reduced Matrigel Matrix basement membrane (REF 356231, lot 7114006, Corning). For passaging the cells Accutase (StemCell Technologies, Vancouver, BC) was used for 5 min at 37C and then stopped and pelleted by gentle centrifugation with iPSC media containing 10μM Y-27632 (Selleckchem, Houston, TX) (Bajpai et al., 2008). The dissociated iPSCs were seeded in new plates with media containing Y-27632, and after 24 hours replaced with media alone. IPSC media was changed daily. Cells were routinely tested and confirmed negative for Mycoplasma contamination by MycoAlert (Lonza).

### Lentiviral constructs and protocol

Lentiviral constructs CS-TRE-c-MYC-PRE-Ubc-tTA-I2G-RESS-GFP (cMYC) and CS-TRE-c-MYC-PRE-Ubc-tTA-I2G-RESS-GFP (BCL-XL) were generous gifts from the Eto laboratory and were transduced as previously reported (Hirose et al., 2013).

### FACS analysis of cells

After lentiviral infection by the Doxycycline-inducible cMYC or Doxycycline-inducible (BCL-XL) vectors, cells were separated on a FACS AriaII for GFP positive cells. Individual clones were isolated by sorting single iPSCs to wells of a 96-well plate precoated with Matrigel with iPSC media containing 10μM Y-27632 and 2μM doxycycline. Clonal iPSC isolates were expanded by weekly passaging and standard iPSC culture as described above.

#### iPSC-Heps differentiation

##### Induction of Definitive Endoderm (days 0-7)

IPSCs were plated on 6, 12, or 96-well Matrigel-coated plates (growth factor reduced, REF 356231, lot 7114006, Corning) at different densities for differentiation. For example typical plating density in a single well of a 6 well plate was 500,000 cells or 20,000 cells per well for 96 well plates. The differentiation process consisted on one week of RPMI media supplemented with Gem21 NeuroPlex without insulin (Gemini Bio-Products catalog #400-962, Gemini, West Sacramento, California), Glutagro catalog# 25-015-Cl (Corning, Manassas, VA), Non-essential amino acids catalog # 11-140-050 (Gibco), Sodium Butyrate (0.5 mM) catalog # B5887 (Sigma Aldrich) and recombinant human Activin A (100 ng/mL) Peprotech catalog #120-14E for 7 days at 20% oxygen, 5% C0_2_. On the day before differentiation (day 0), cells were re-plated on Matrigel coated plates in the presence of 2μM doxycycline and 10μM Y-27632. On day 1 of induction, 2% KnockOut Serum Replacement (KSR) Gibco Catalog # 10828028, CHIR-99021 Selleckchem catalog # S2924 (3mM), PI-103 (50nM) Selleckchem catalog # S1038 and recombinant human BMP4 (10ng/mL) Peprotech catalog #120-05ET and FGF2 (20ng/mL) Peprotech catalog #100-18B were added to the media. On day 2, 1% KSR, PI-103 (50nM) and BMP4 (10ng/mL) and FGF2 (20ng/mL) and on day 3, 0.2% KSR and PI-103 (50nM) were added to the media. Doxycycline hyclate (Acros Organics catalog #446061000) was added throughout definitive endoderm induction at various doses as described in text with a standard dose being 2μM.

##### Differentiation of Definitive Endoderm to hepatocytes (days 8-23)

IPSC-derived human endoderm from above was cultured continuously on Matrigel or split up to 1:4 onto Matrigel plates for hepatic differentiation. Definitive endoderm (DE) was cultured in Iscove’s modified Dulbecco’s medium (Gibco catalog # supplemented with Gem21 NeuroPlex without insulin (Gemini Bio-Products catalog #400-962), Glutagro, NEAA, 0.3 mM monothioglycerol, 0.126 M/mL human insulin (Sigma), and 100 nM dexamethasone. To induce hepatoblasts, cells were treated with FGF2 (10 ng/mL) and BMP4 (20 ng/mL) for 5 days at 5% oxygen, 5% C0_2_. To further differentiate the cells towards hepatocytes, cells were treated with FGF2 (10 ng/mL), BMP4 (20 ng/mL), and recombinant human HGF (20 ng/mL) Peprotech catalog #100-39 for 5 days at 5% oxygen, 5% C0_2_. This was followed by changing the culture conditions to Lonza Hepatocyte Culture Media BulletKit (HCM Catalog # CC-3198) with HGF (20 ng/mL) and recombinant human Oncostatin M (20 ng/mL) Peprotech catalog # 300-10 for 5 additional days at 20% oxygen, 5% C0_2_. In the BulletKit, use of EGF was excluded.

##### Immunohistochemistry and Cell staining

IPSCs and iHeps were fixed using 4% paraformaldehyde for 15 min, permeabilized using 0.1% triton X-100 for 10 minutes followed by 3x PBS (phosphate buffered saline, Corning Life Sciences) rinses and blocked with 1.5% bovine serum albumin for 1 h. Samples were incubated with primary antibody overnight (List of all antibodies with dilutions 1:100 - 1:., OCT3/4 (Santa Cruz, #sc-9081, 1:500, SOX2 (Proteintech, #11064-1-AP, 1:1000), NANOG (Proteintech, #14295-1-AP, 1:1000), SSEA (Proteintech, #19497-1-AP, 1:500), TRA 1-60 (Proteintech, #18150-1-AP, 1:500), HNFα4 (Proteintech, #26245-1-AP, 1:500, Albumin (Bethyl, #A80-129A, 1:500, AFP (Thermo Scientific, #RB-365-A1, 1:500), ki-67 (8D5) (Cell Signaling, #9449, 1:800), Cleaved Caspase-3 (Cell Signaling, #9661, 1:400)) in blocking solution. The cells were rinsed extensively with PBS and incubated for 60 min with secondary antibody at 1:1000 in blocking solution. The secondary antibodies used were: Alexa Fluor 555 donkey anti-mouse IgG (H+L), Alexa Fluor 488 goat anti-rabbit (H+L) (Invitrogen, Carlsbad, CA). After washing 3 times with PBS, DAPI (4’,6-diamidino-2-phenylindole, MP Biomedicals, Solon, OH) was added at 300 nM for 15 min. After 3x PBS washes, the cells were imaged using an BIOTEK Cytation 5 Microscope (Winooski, VT). MitoTracker Orange CMTMRos staining was performed as follows: iPSCs were plated, induced 24 hours with or without 1mM doxycycline, and 24 hours later, Mitotracker Orange dye was added for 45 minutes in PBS buffer at 37 degrees for 30 minutes, and then rinsed in PBS. Cells were then fixed with 4% PFA and Hoechst 33342, rinsed in PBS and imaged with Cytation5 imager, BioTek Instruments. Image were analyzed using Biotek Imaging software. Primary and secondary mask analyses on digital images were used to quantify based on nuclei and antibody staining in secondary mask. Threshold analysis was set to exclude artifacts.

##### RNA isolation, Real-Time PCR analysis, and RNA Sequencing

Total RNA was isolated by Qiagen RNeasy mini kit or Zymo Direct-zol RNA kits. Complementary DNA (cDNA) was synthesized by qScript cDNA SuperMix (Quantabio). Quantitative Real-Time PCR (qrtPCR) was performed using FastStart Universal SYBR Green, Roche Diagnostics (Indianapolis, IN) Primers were made by IDT. Primers used for qrtPCR include cMyc-F 5’-AAACACAAACTTGAACAGCTAC-3’, cMyc-R 5’-ATTTGAGGCAGTTTACATTATGG-3’, BCL2L1-F 5’-TCCTTGTCTACGCTTTCCACG-3’, BCL2L1-R 5’-GGTCGCATTGTGGCCTTT-3’, GAPDH-F 5’-AGCCACATCGCTCAGACAC-3’, GAPDH-R 5’-GCCCAATACGACCAAATCC-3’. RNA Sequencing was performed by BGI, Boston, MA. Poly-A RNA was selected using oligo-dT magnetic beads followed by N6 random priming. On average 27,642,407 raw reads were obtained and mapped per sample. Raw reads were subjected to quality control. Clean reads were analyzed by Gene Ontology analysis, KEGG pathway enrichment and cluster analysis. RNA Sequencing data was deposited and is available at ArrayExpress accession number E-MTAB-8821.

##### Moon Bio Antibody Array

To analyze protein levels and specific phosphorylation, the Phospho Explorer Antibody Array was performed. IPSC line 7192 was induced with and without doxycycline 1μM for 3 days. Extracts were made using Complete Lysis-M EDTA-Free Lysis buffer (Roche). Antibody Array was performed, scanned, and analyzed by Full Moon Biosystems, Sunnyvale, CA.

##### Western blot analysis

Performed as previously described (Mattis et al., 2015). Protein bands were visualized by Odyssey Infrared Imager scanner and band intensities were quantified using the Image Studio software (LI-COR Biosciences).

#### Statistical Analysis

Experiments were run in at least triplicate for each condition. Statistical significance was represented as standard error of the mean (SEM) and was determined using Student t test or Anova analysis as appropriate (GraphPad Prism, La Jolla, CA).

#### Study Approval

All studies using human stem cell materials were carried out under approval by the University of California, San Francisco, Institutional Review Board (IRB) study approval 10-04393.

## Author Contributions

C.E.F. and T.S. carried out experiments for Figures 1–5. C.P. and C.D. performed additional differentiation experiments on multiple cells lines to confirm the method was working and performed experiments for supplementary figures. K.L. and M.M. provided bioinformatics analysis. J.J.M. provided critical guidance over western blots and repeated experiments in her own lab on novel set of iPSCs. A.N.M. conceived of the experiments, made the figures, and wrote the paper.

## Acknowledgements

This study was supported in part by National Institutes of Health (NIH) Grant K08DK098270 to ANM, the UCSF Department of Pathology and the UCSF Liver Center (P30 DK026743).

---

**Supplementary Figure 1.**
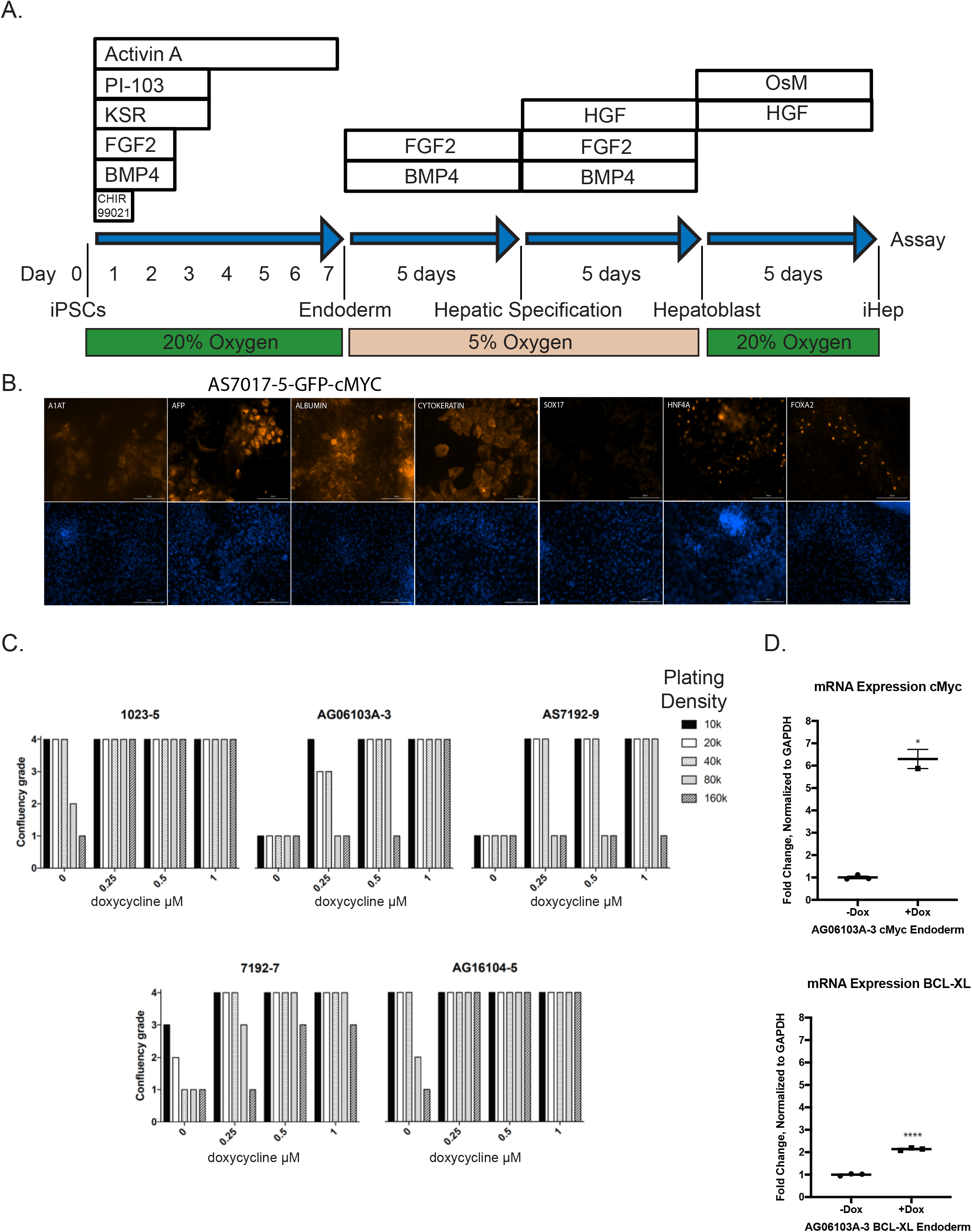
Standard differentiation protocol. (A) Our old protocol for induction of iPSC to endoderm and iPSC-Heps. (B) Induction of iPSC with doxycycline inducible cMYC results in iPSC-Heps with appropriate staining of markers including A1AT, AFP, Albumin, HNF4α and FOXA2. (C) Plating density versus confluency after 6 days of induction (n=2 technical replicates in 5 different cell lines, line AS7192 was tested in two different subclones designated −9 and −7). Confluency grade analyzed visually and defined as 1-no cells remain, 2-few cells, 3-about 75% covered well, 4-complete monolayer. (D) Quantitative PCR for mRNA expression of cMYC relative to GAPDH in lentivirally transduced AG06103A-3 with and without doxycycline induction (n=3). Data were analyzed by Student’s t test and presented as mean ± SEM. * p < 0.05; ** p < 0.01.

**Supplementary Figure 2.**
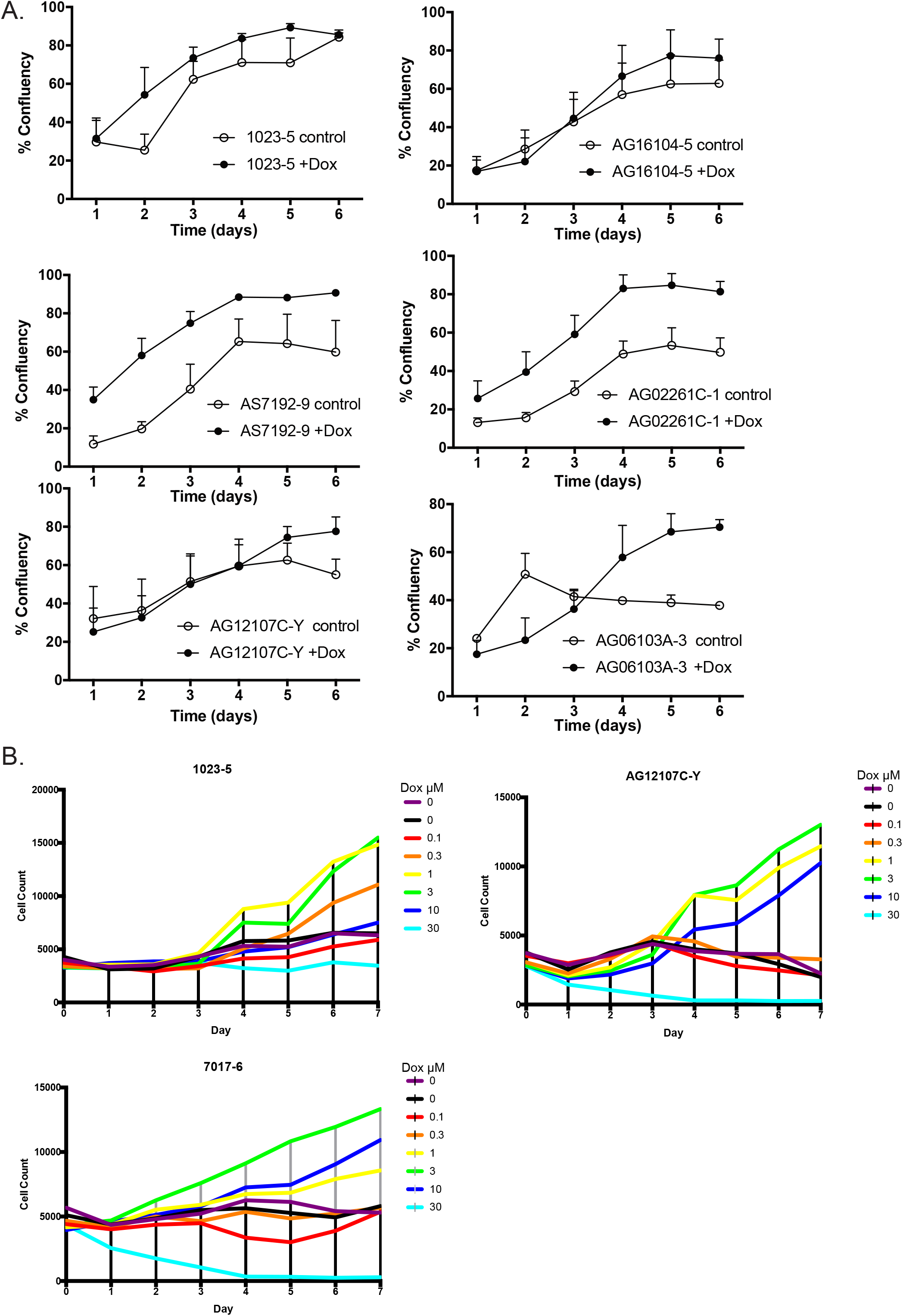
Induction with Doxycycline and Dose Response. (A) Comparison of control versus 1μM doxycycline (+Dox) induction over six different cell lines shows improved confluence and survival of cells (n=3). (B) Titration of Doxycycline dosage shows that there is an optimal level of doxycycline and a toxic level (n=3 biologic replicates – performed once in three different cell lines). Data were analyzed by Student’s t test, error bars represent 95% confidence interval.

**Supplementary Figure 3.**
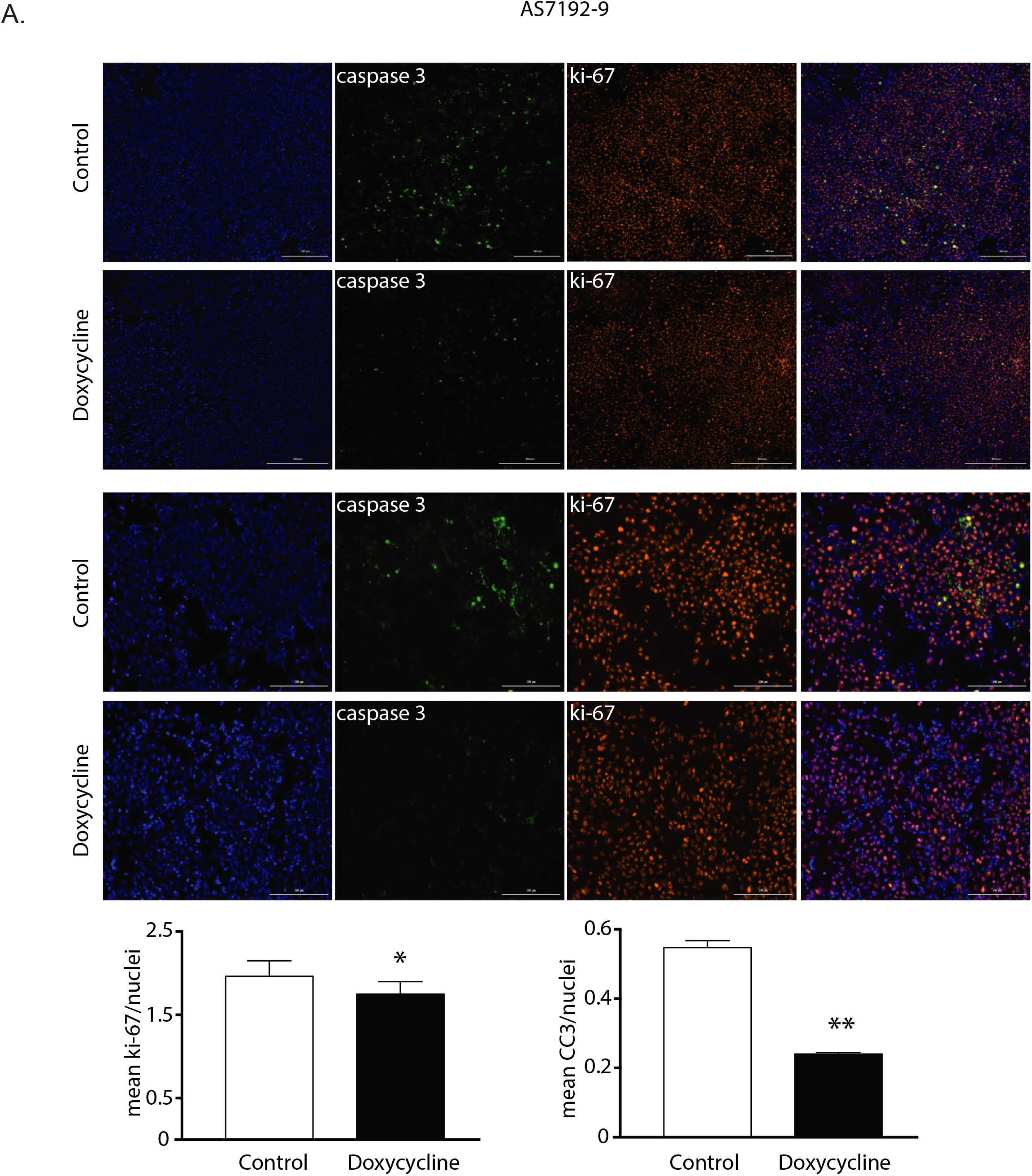
Doxycycline Reduces Cleaved Caspase 3 Levels. (A) Immunofluorescence of iPSC line AS7192-9 two days after induction with staining for Hoechst 33342, Ki-67, Cleaved Caspase 3 (CC3), and combined (first column) shows greatly decreased CC3 in the presence of doxycycline (n=3). At this time point, there is a slight decrease in Ki-67. First two rows are 10x with scale bar at 400μm and bottom two rows 20x with 200μm scale bar. Data were analyzed by Student’s t test and presented as mean ± SEM. * p < 0.05; ** p < 0.01.

**Supplementary Figure 4.**
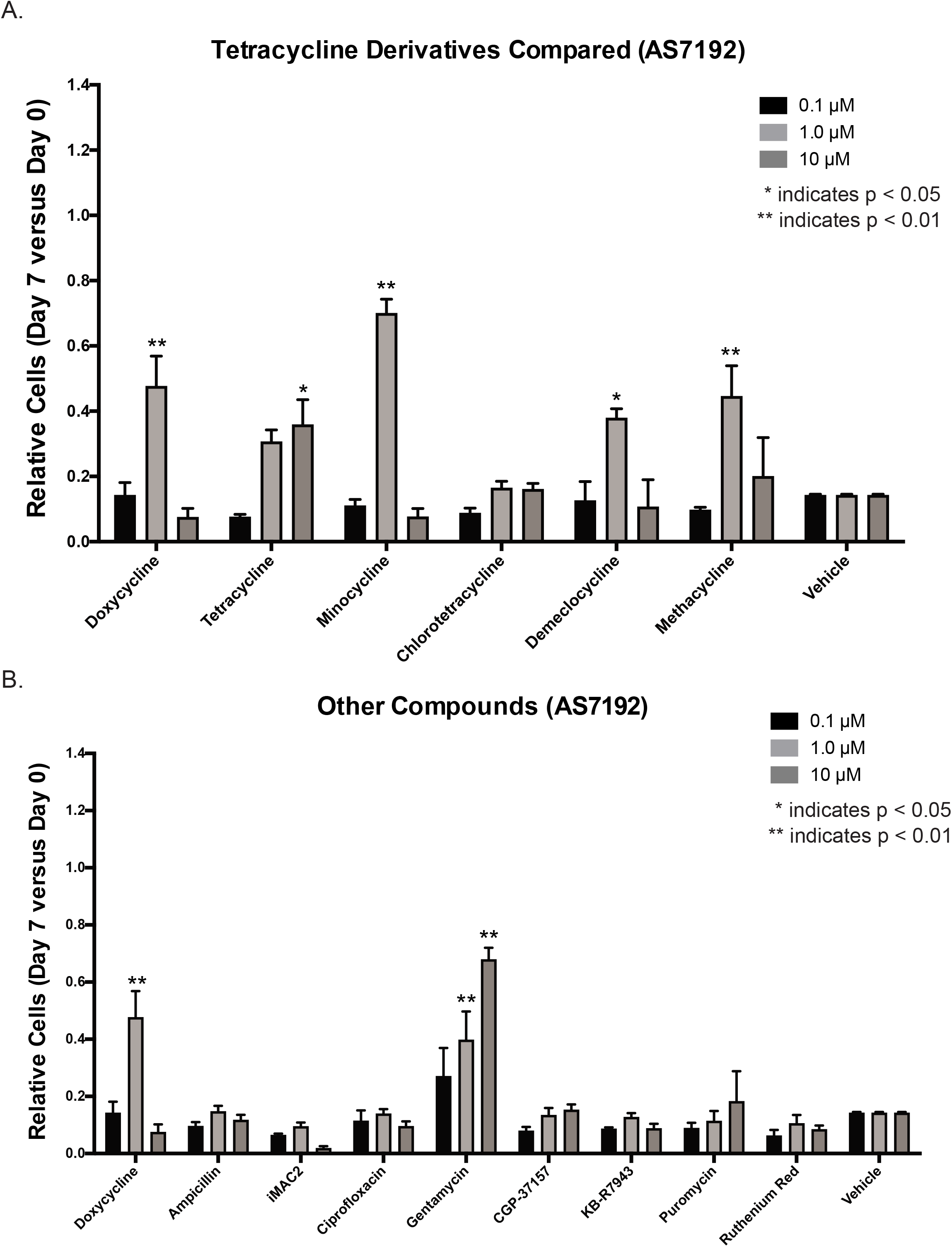
Comparison of Tetracycline Derivatives and other compounds in iPSC AS7192. (A) Induction of iPSC cell lines AS7192 to endoderm over 7 days was performed. Doxycycline was compared to tetracycline derivatives in its ability to rescue endoderm induction. Minocycline, and Methacycline showed similar ability to rescue endoderm at the 1 μM concentration with some toxicity at the 10μM concentration. Demeclocycline showed a similar trend. The number of cells is shown measured seven days after induction compared to the plating density at day 0 (n=3). (B) Other antibiotics and other compounds affecting the mitochondria were compared to doxycycline. While most antibiotics failed to have the same affect, surprisingly Gentamycin was shown to have a strong increasing dose response. The number of cells is shown measured seven days after induction compared to the plating density at day 0 (n=3). Data analyzed by two way ANOVA and presented as mean ± SEM. * p < 0.05; ** p < 0.01.

**Supplementary Figure 5.**
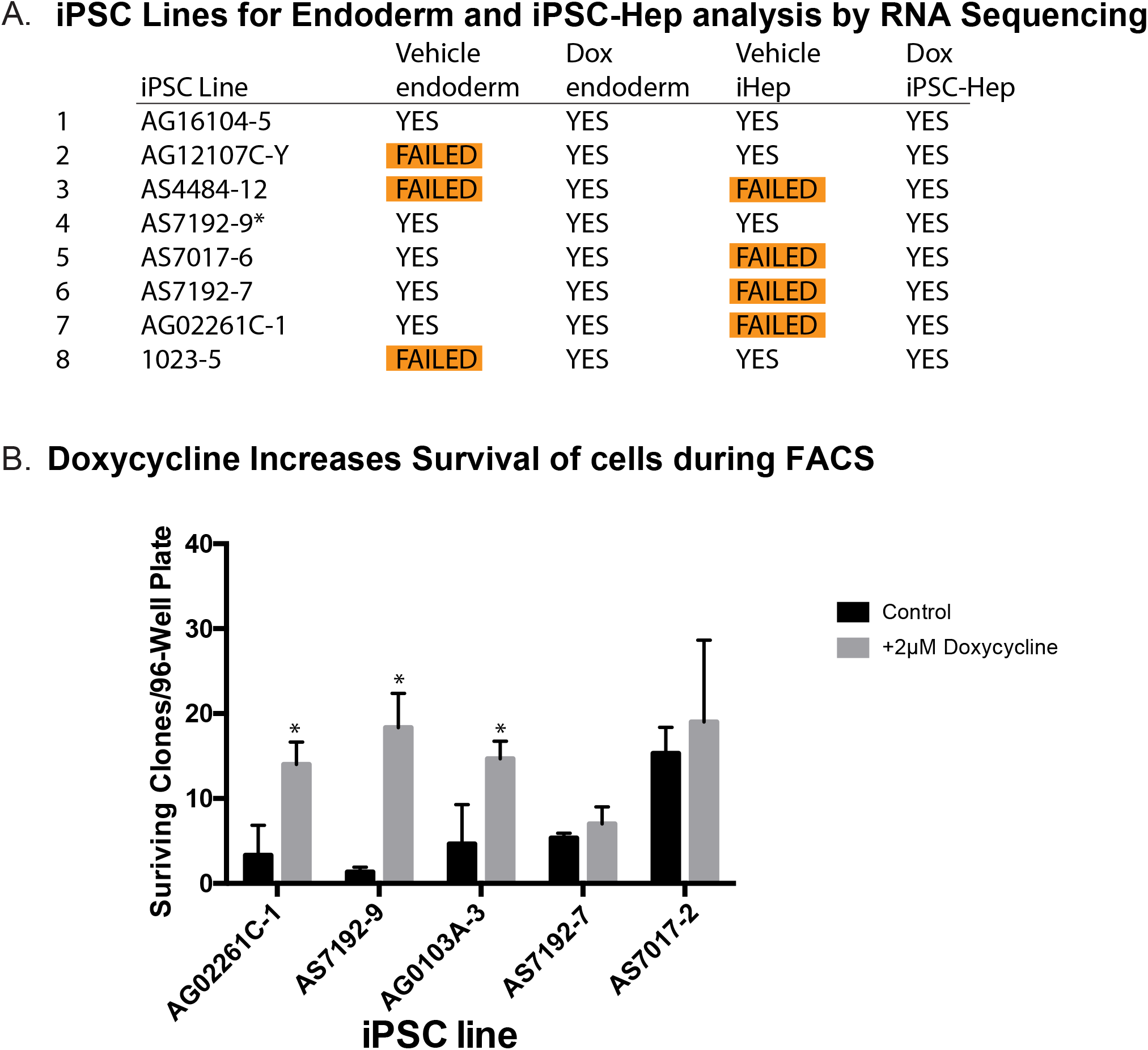
Induction of Lines for RNA Sequencing and effects of Doxycycline on FACS. (A) iPSC Lines differentiated to Endoderm and iPSC-Hep for analysis by RNA Sequencing. Without doxycycline a number of cell lines underwent apoptosis before successful endoderm or did not produce enough cells at the iPSC-Hep stage. Two separate differentiations of AS7192-9 were produced at the iPSC-Hep stage (designated by *). Some iPSC lines failed validation for RNA Sequencing and were excluded in the RNA Sequencing data. Additional controls were also used for RNA Sequencing designated controls 1 through 4. (B) Doxycycline (2μM) increases the survival of cells during FACS sorting especially for sorting of single cells to create clonal lines (n=5). Data were analyzed by two way ANOVA and presented as mean ± SEM. * p < 0.05; ** p < 0.01.

